# The potential effects of tree planting on allergenic pollen production in New York City

**DOI:** 10.1101/2023.04.11.536416

**Authors:** Daniel S.W. Katz

## Abstract

Tree selection decisions affect urban pollen production but the overall importance of tree planting to airborne pollen concentrations remains poorly understood. A synthesis of existing data and available literature could inform tree -planting decisions and potentially help reduce future airborne pollen concentrations. This is especially relevant for the many cities that are considering tree planting campaigns, such as New York City. Here, I examine which allergenically -important tree taxa could be most influenced by municipal tree selection decisions in New York City by comparing tree species abundance from a representative plot-based city-wide tree survey with a street tree inventory. I then estimate pollen production from several allergenic tree taxa by combining these tree datasets with allometric equations of pollen production as a function of tree size. Pollen production is also compared to several years of airborne pollen measurements. The potential effect of a proposed planting campaign is estimated over time by combining growth rate equations with pollen production equations. Several tree genera are especially important producers of allergenic pollen in New York City, including *Quercus, Platanus, Morus*, and *Betula*; these taxa also comprise 71% of airborne pollen measured and 93% of estimated pollen production (107 quadrillion pollen grains ; however pollen production could not be estimated for all taxa). *Platanus* × *acerifolia* is predominantly a street tree, indicating that previous municipal planting decisions have resulted in its current abundance (it accounts for 34% of total street tree basal area and has an estimated annual pollen production of almost 30 quadrillion grains) and will determine its future abundance. In contrast, *Morus* and *Betula* are uncommon as street trees, indicating that municipal tree planting campaigns are unlikely to substantially affect their pollen production rates in NYC. *Quercus* was the largest estimated producer of pollen in NYC (62 quadrillion pollen grains) and accounted for almost 25% of airborne pollen collected, but its very high abundance outside of street trees suggest that the relative effect of planting trees in this genus will be relatively small. Overall, this study demonstrates how tree planting decisions can have important and long-lasting consequences for allergenic pollen production in certain circumstances, suggesting that pollen allergenicity should be considered in future tree selection decisions.

## Introduction

Exposure to pollen results in allergic reactions that substantially reduce quality of life for the millions of Americans that are sensitized to pollen (Meltzer 2016; Meltzer et al. 2009). Urban trees produce large quantities of pollen (Katz et al. 2020; Maya-Manzano et al. 2017) and tree pollen has been implicated in elevated rates of asthma -related emergency department visits in spring (Anenberg et al. 2017; Darrow et al. 2012; Sun et al. 2016). There have been several calls for selecting plants with low allergenic potential (Cariñanos and Casares-Porcel 2011; Green et al. 2018; Maya-Manzano et al. 2017; D. J. Nowak and Ogren 2021), but relatively little is known about the potential for tree planting decisions to affect pollen production, airborne pollen concentrations, pollen exposures, and the resulting health consequences including allergic rhinitis symptoms and asthma exacerbations. A synthesis of relevant information and datasets is needed to link tree selection to pollen production and the ensuing public health consequences; a better understanding of these relationships could inform management decisions, such as tree planting campaigns.

The allergic consequences of tree planting decisions depend in part on the proportion of trees that are intentionally planted compared to those that are naturally recruited (D. J. Nowak 2012; D. J. Nowak and Ogren 2021). Specifically, tree selection decisions can gradually shift the composition of planted trees but have weaker effects on natural areas, vacant lots, and other minimally managed areas (D. J. Nowak 2012). Street trees, which often account for ∼5-10% of total trees in cities in the United States (McPherson et al. 1997, 2016), are owned by cities and their composition is in part the result of centralized decisions. Thus, from a municipal government perspective, allergenic pollen production from street trees is both a responsibility and a problem that could be addressed with little extra effort when planting new trees.

The long lifespan of trees means that t ree planting decisions have long-term consequences; current pollen production is often the result of decisions made several decades ago. The exact timescales of these effects will depend upon tree growth and mortality rates which vary among taxa, regions, and urbanization gradients (McPherson et al. 2016; Rötzer et al. 2020). Certain species may also display non - linear increases in pollen production as a function of tree size (Katz et al. 2020) or may reach sexual maturity at varying ages or sizes. Thus, the time-scales over which planting decisions will affect pollen production is an important consideration, but little work has quantified changes in tree pollen production over time due to tree growth and aging.

Here, I explore the potential for tree planting decisions to affect pollen production using New York City (NYC) as a case study. Specifically, I quantify current pollen production by street trees compared to city-wide tree composition and compare tree abundance to airborne pollen measurements. I then assess how current tree selection decisions could affect pollen production over time. Finally, I review and discuss various relevant processes, including the spatial scales of pollen production and dispersion, the role of atmospheric dispersion, botanical sexism, and generalizability to other metropolitan areas.

## Methods

### Study site description

New York City is home to 8.5 million people; 11% of children there have asthma (New York State Department of Health 2009). Asthma is closely connected to allergic rhinitis (Giavina-Bianchi et al. 2016; Grossman 1997), and pollen is one of the major triggers of seasonal allergies (Bousquet et al. 2008). In NYC, airborne pollen concentrations have been linked to asthma-related emergency department visits in the Bronx (Jariwala et al. 2011, 2014) and tree cover has been linked to pollen sensitization (Lovasi et al. 2013).

New York City is currently 22% forested (17,254 ha or 42,635 acres), with approximately half of that in areas managed by the NYC Department of Parks and Recreation, including 28% in parks (approximately one third of which is in landscaped areas) and 25% in rights of way including street trees; about 35% is on privately owned land (Maxwell et al. 2021a). From 2010 to 2017, at least 518 ha of new canopy developed separately from existing tree canopy; this can coarsely be assumed to be the result of new plantings, such as the 136,871 street trees and 17,440 trees planted in landscaped park areas during that period (Maxwell et al. 2021b). A recent publication concluded that tree cover in NYC could reasonably be increased from 22% to between 30 – 42% (Treglia et al. 2022). Further tree planting campaigns are gaining momentum in NYC, as exemplified by the *Million More Trees* proposal (Rubinstein 2022). Tree planting campaigns are also gaining ground nationally (Eisenman et al. 2021a; Sousa-Silva et al. 2023) and the Inflation Reduction Act of 2022 will invest an additional $1.5 Billion in urban and community forestry.

### Data sources

To investigate the potential for tree planting to affect pollen production in NYC, I compared species composition and pollen production across the city with that of street trees. One of the primary sources of data on trees across NYC was a plot-based survey in the summer of 2013 that sampled 296 plots, each of which was randomly placed and covered one tenth of an acre (D. Nowak et al. 2018). This sampling program used the i -Tree Eco sampling approach and counted and identified all woods plants with a diameter at breast height (DBH) > 1 inch; size class of each tree was also recorded. In total, 1,075 trees were measured.

As with many cities, the majority of mapped trees are street trees (Keller and Konijnendijk 2012). In 2015 and 2016, a street tree inventory was implemented by NYC Parks staff and volunteers (NYC Parks Department 2017). In total, 652,088 living street trees were identified and mapped; these data are publicly available at: https://tree-map.nycgovparks.org/. DBH was also recorded for each tree.

Airborne pollen data were collected in Manhattan using a Burkhard sampler on the Lincoln Center roof. Pollen was collected during the main pollen seasons of 2009 – 2012; the first day of collection in most years was March 1 (with the exception of 2009, where it was April 6), and the final day of collection was Oct 31. Pollen was identified to 87 taxonomic groups; these were generally at the genus level, with some exceptions where identification is only feasible at the family level (e.g., Poaceae/Gramineae, Cupressaceae, and Pinaceae) or where pollen types cannot be readily distinguished (e.g., *Carpinus* and *Ostrya*). Data were generously provided by Dr. Guy Robinson of Fordham University.

### Data analysis

Pollen production was estimated as a function of tree size for several species using the equations developed by Katz et al., 2020a. Briefly, these equations were based on pollen production estimates of urban trees for 10 tree species in Ann Arbor, Michigan; for each species pollen production of at least 10 individuals were measured; among species the average R ^2^ = 0.72 (range of R^2^: 0.41 – 0.99). Pollen production equations were available for the following species: *Acer negundo, Acer platanoides, Acer rubrum, Acer saccharinum, Betula papyrifera, Gleditsia triacanthos, Juglans nigra, Morus alba, Platanus* × *acerifolia, Populus deltoides, Quercus palustris, Quercus rubra*, and *Ulmus americana*.

Within *Betula, Morus, Quercus*, and *Ulmus*, I applied the same pollen production equations to other species within that genus, including: *Betula lenta, Betula nigra, Betula occidentalis, Betula populifolia*, unidentified species of *Morus, Quercus alba, Quercus bicolor, Quercus phellos, Quercus prinus, Quercus velutina, Ulmus parvifolia*, and unidentified *Ulmus* species. Due to the high variation in tree structure, pollen production, and pollination within *Acer*, pollen production was not extrapolated to other congeneric species. City-wide pollen production was calculated by summing total pollen production for each tree in each size class of the included genera from the i -Tree data. Street tree pollen production was estimated based on the size (DBH) of each tree. These estimates adjust for tree sex for diecious *Morus* (male: 57.8%) and for the polygamodioecious *Acer rubrum* (male: 10.6%) based on observed sex ratios presented elsewhere (Katz et al. 2020). For taxa that had non-linear relationships between pollen production and basal area (*Morus, Populus*, and *Ulmus*), total calculated pollen production was limited to that of the observed maximum tree size from the original dataset used to create the pollen production estimates, to prevent potentially unrealistic pollen estimates for very large trees.

To calculate the effect of tree planting on pollen production, we combined tree growth rate equations (McPherson et al. 2016) and pollen production equations (Katz et al. 2020) to estimate first DBH as a function of age and then pollen production as a function of basal area (derived from DBH). This allowed us to both estimate the effects over time. To create this simulation, we assumed a p lanting campaign of 200,000 trees, which is the estimated capacity for additional street trees in NYC (Treglia et al. 2022). Proposed planted trees were assumed to be proportional to existing street tree composition (i.e., tree taxonomic identity was sampled with replacement from the street tree database). Pollen production was then calculated over 50 years based on increased in growth rate. Neither mortality nor changes in currently existing trees were included in this simulation.

Data were analyzed in R 4.1.2 (R Core Team 2018) using several packages within the Tidyverse (Wickham et al. 2019). Upon publication, code will be available at: https://github.com/dankatz/nyc_tree_planting.git

## Results

Several tree genera of allergenic concern are both prevalent and achieve high airborne pollen concentrations in NYC, including *Quercus, Platanus, Betula*, and *Morus* (Table 1; a version of Table 1 including all taxa is available in SI Table 1). Together, these taxa account for 71% of the airborne pollen monitored at the Lincoln Center pollen monitoring station. Of particular note is the strong disjunct between relative basal area and share of airborne pollen for certain taxa; for example, *Betula* is estimated to include 1.7% of total basal area in NYC but accounts for 12% of measured airborne pollen. Similarly, *Morus* accounts for 0.7% of total basal area but 23% of airborne pollen. In contrast, *Quercus* accounts for 27% of total basal area and 24% of airborne pollen. With a basal area of 8% and accounting for 12% of airborne pollen, *Platanus* is an important producer of allergenic pollen and it is especially well represented among street trees, accounting for 34% of all street tree basal area. Table SI 2 summarizes the street tree database by species and Table SI 3 summarizes the i-Tree data by species.

**Table 1.** Allergenic pollen producing trees in New York City based on citywide i-Tree plots, street tree database, and airborne pollen collection. Any taxon with >1% relative basal area (in either the i -Tree plots or the street tree database) or with >1% proportion of total pollen collected is included here. *Pollen concentrations provided here for *Picea* also include other members of the Pinaceae family, including *Pinus*. **Pollen concentrations provided for *Thuja* also include other members of the Cupressaceae family, including *Juniperus, Cryptomeria*, and *Taxodium*. For wind pollination, + = generally wind pollinated, - = few wind-pollinated species. For allergic concern + = of allergic concern, - = not of allergic concern, and ∼ = of unknown or minor allergic concern.

| taxon | i-Tree relative basal area (%) | i-Tree relative number of stems (%) | Street tree relative basal area (%) | Street tree relative number of stems (%) | Pollen (% collected) | Pollen max (grains/m<sup>3</sup>) | Wind pollinated | Of allergenic concern |
| --- | --- | --- | --- | --- | --- | --- | --- | --- |
| *Quercus* | 26.52 | 10.52 | 18.21 | 12.71 | 23.9 | 2158.8 | + | + |
| *Acer* | 19.68 | 11.78 | 15.57 | 13.61 | 1.0 | 119.3 | + | + |
| *Platanus* | 8.29 | 2.38 | 34.47 | 13.34 | 11.9 | 2727.6 | + | + |
| *Liquidambar* | 3.49 | 1.58 | 1.38 | 1.63 | 0.5 | 87.2 | + | ~ |
| *Tilia* | 3.22 | 0.96 | 5.05 | 7.86 | <0.1 | 11.0 | - | ~ |
| *Robinia* | 3.16 | 4.08 | 0.30 | 0.27 | <0.1 | 4.0 | - | - |
| *Ulmus* | 2.15 | 1.19 | 1.95 | 2.29 | 1.1 | 176.8 | + | + |
| *Crataegus* | 2.11 | 1.25 | 0.05 | 0.51 | 0.0 | 0.0 | - | - |
| *Fagus* | 2.04 | 0.26 | 0.02 | 0.06 | 0.1 | 27.0 | + | - |
| *Fraxinus* | 1.78 | 0.57 | 3.12 | 3.06 | 2.0 | 437.5 | + | + |
| *Picea* | 1.75 | 1.09 | 0.05 | 0.10 | 4.2* | 365.1* | + | - |
| *Betula* | 1.66 | 4.87 | 0.05 | 0.21 | 12.3 | 2396.8 | + | + |
| *Pyrus* | 1.59 | 2.49 | 4.39 | 9.04 | 0.0 | 0.0 | - | - |
| *Tsuga* | 1.58 | 0.82 | 0.00 | 0.01 | 0.1 | 33.1 | + | - |
| *Gleditsia* | 1.52 | 0.92 | 6.11 | 9.85 | 0.3 | 130.4 | + | ~ |
| *Celtis* | 1.5 | 2.18 | 0.10 | 0.37 | <0.1 | 2.2 | + | + |
| Unknown | 1.42 | 3.4 | 0.00 | 0.00 | 3.1 | 189.5 |  |  |
| *Sophora* | 1.41 | 0.46 | 0.00 | 0.00 | 0.0 | 0.0 | - | - |
| *Thuja* | 1.1 | 6.32 | 0.01 | 0.05 | 6.4** | 1076.1** | + | + |
| *Ailanthus* | 1.02 | 5.54 | 0.13 | 0.12 | NA | NA | - | ~ |
| *Prunus* | 0.93 | 3.35 | 1.37 | 6.39 | <0.1 | 43.7 | - | - |
| *Zelkova* | 0.71 | 0.72 | 1.91 | 4.49 | 0.2 | 295.63 | + | + |
| *Morus* | 0.69 | 0.82 | 0.29 | 0.18 | 22.6 | 4342.6 | + | + |
| *Ginkgo* | 0.57 | 0.53 | 1.61 | 3.22 | 0.7 | 311.1 | + | ~ |
| *Styphnolobium* | 0.00 | 0.00 | 1.64 | 2.97 | 0.0 | 0.0 | - | - |
| *Populus* | 0.00 | 0.00 | 0.11 | 0.07 | 1.0 | 156.8 | + | + |

Based on i-Tree plot estimates of tree abundance and size and pollen production equations, the genera with the highest estimated pollen production were *Quercus, Platanus, Morus*, and *Betula* (Fig. 1). The single species with the highest estimated pollen production was *Platanus* × *acerifolia*, with a total annual pollen production of 29.6 quadrillion pollen grains in NYC. Pollen production equations were not available for species in the Cupressaceae and Pinaceae families, so pollen production estimates for those taxa are not included here; nor are equations for other species of *Acer*. For street trees, the genera with the highest estimated pollen production were *Platanus, Quercus, Gleditsia*, and *Ulmus* (Fig. 1). *Platanus* were the largest street trees; their average DBH was 54.7 cm, substantially greater than any other species and there were relatively few small *Platanus* trees.

**Fig. 1:**
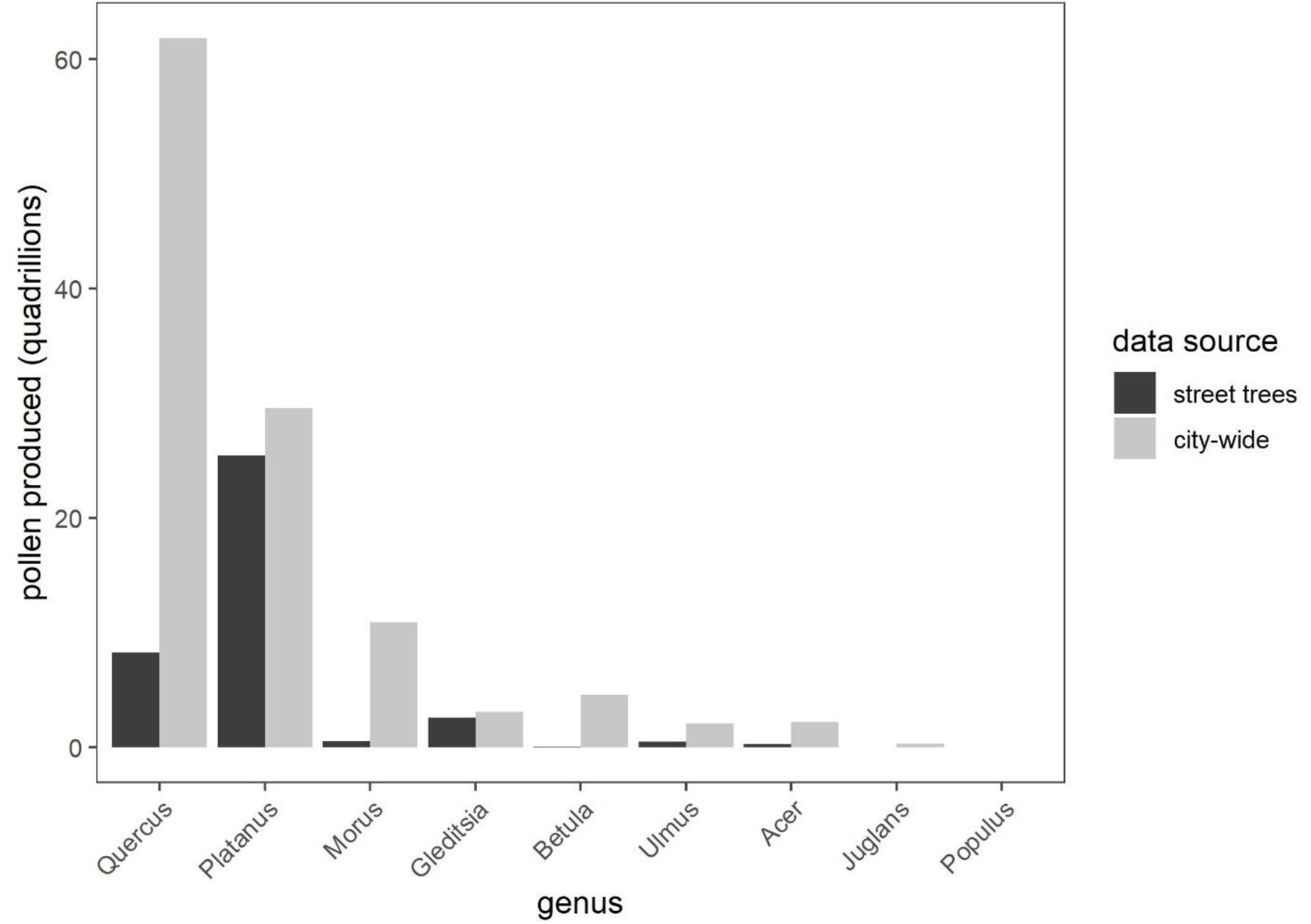
Estimated total pollen production in quadrillions of pollen grains for tree genera for all trees across NYC and for street trees. Pollen production is based on abundance in each size class for taxa for which allometric equations are available. Citywide data comes from i -Tree plot estimates.

The effects of simulated tree planting on pollen production were low for most species when tree planting was representative of current street trees (Fig. 2). In some cases, this is due to low representation of particular allergenic pollen producing trees amongst street trees. For example, in the 2015 street tree census there were only 1,156 *Morus* street trees (0.18% of total street trees) and 1,400 *Betula* trees (0.21% of total street trees). For other taxa, the increases in pollen from tree planting were, while large in absolute terms, relatively small compared to existing production. For example, planting an additional ∼25,000 *Quercus* trees is estimated to result in an additional annual production of 8 quadrillion pollen grains 50 years after planting, but this is relatively minor compared to the existing pollen production of 62 quadrillion pollen grains. The taxa for which increases were larger were *Ulmus*, which was expected to increase by approximately 0.5 quadrillion pollen grains over that period (24%), and for *Gleditsia triacanthos*, which was expected to increase by an additional 2.5 quadrillion pollen grains (83%).

**Fig. 2:**
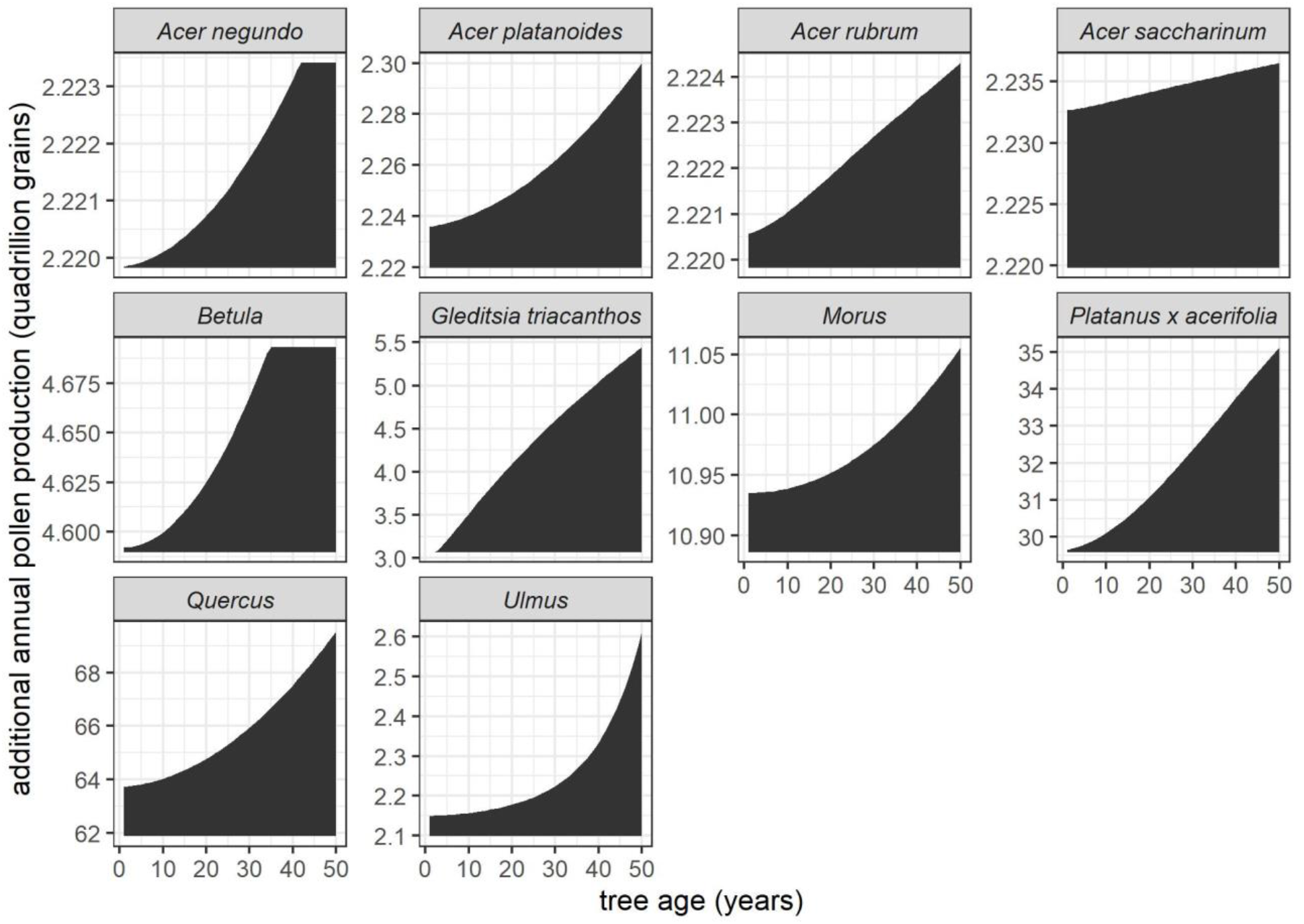
Simulated additional pollen production if an additional 200,000 street trees were planted tha t reflect current street tree composition. Estimated pollen production over time is based on equations for tree size as a function of age and equations of pollen production as a function of size. The y -axis begins at the current city-wide estimate of pollen production for each taxon. Immediate jumps are an artifact of the tree size as a function of age equations, which sometimes start with positive intercepts (i.e., a DBH > 0 at age 0); negative intercepts were restricted to 0. *Betula* size rates were truncated to the range of the data that they were developed from (i.e., we assumed that *Betula* trees could not continue growing forever).

Airborne pollen was collected on a total of 901 days throughout the primary pollen season in 2009 – 2012. The taxa with the highest relative pollen abundance were *Quercus* (24%), *Morus* (23%), *Betula* (12%), *Platanus* (12%), and Cupressaceae (6%); herbaceous taxa including Poaceae, *Ambrosia*, and *Artemisia*, were of far lower importance (Fig. 3). The taxa with the maximum recorded pollen concentrations (Table 1) were *Morus* (4,343 grains/m^3^), *Platanus* (2,728 grains/m^3^), *Betula* (2,397 grains/m^3^), and *Quercus* (2,159 grains/m^3^).

**Fig. 3:**
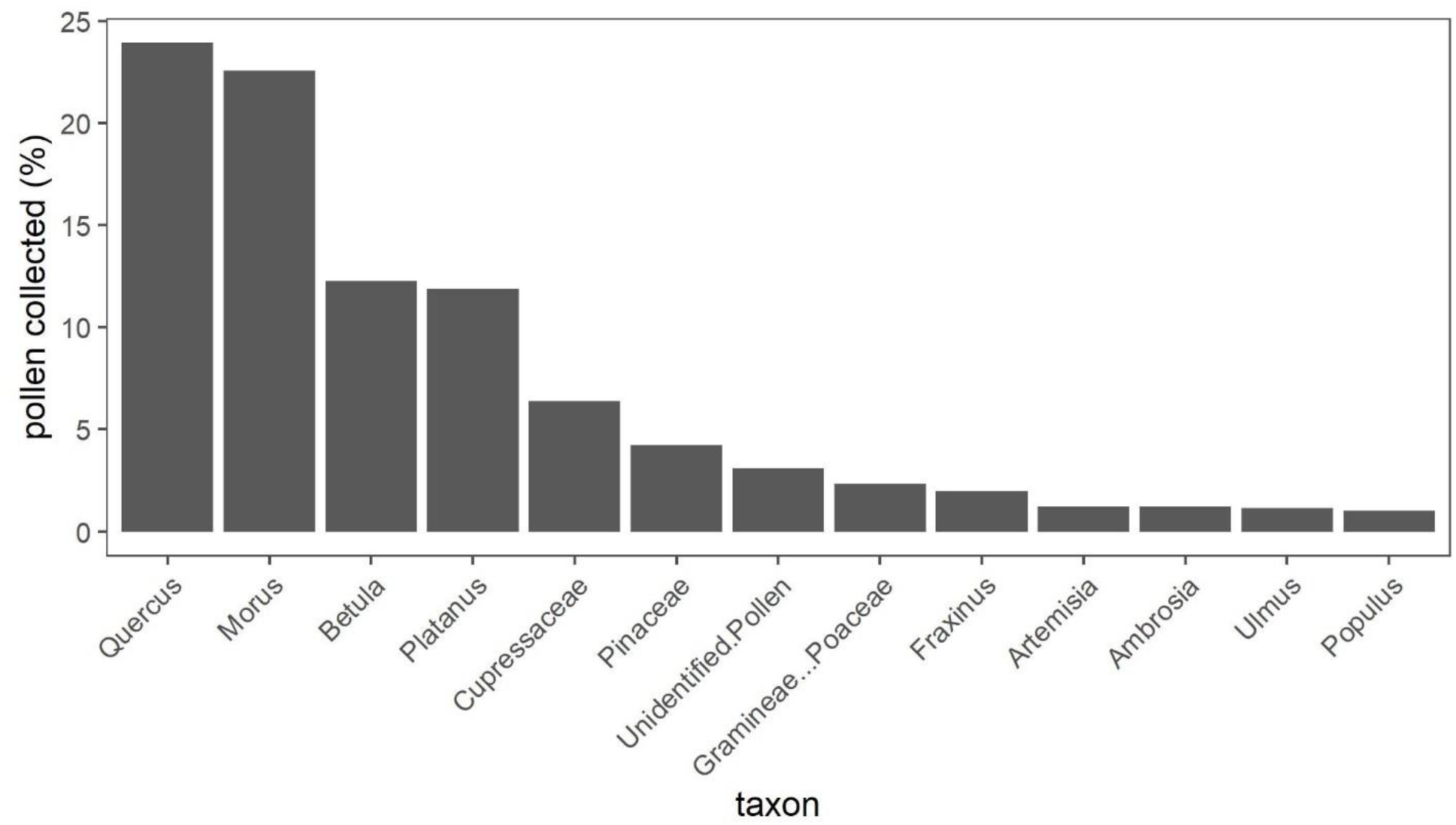
Percent of pollen collected for each taxonomic group for taxa that contributed >5% of total measured pollen. Pollen collection occurred over four years (2009 – 2012) using a Burkard sampler located in Manhattan; data are courtesy of Dr. Guy Robinson.

## Discussion

### Which taxa are of the highest allergenic concern in NYC?

The results presented here demonstrate that, in NYC, municipal tree planting is unlikely to have major effects on city -wide pollen production for all but a few taxa of allergenic concern, as the most important taxa are either already unusual as street trees (*Morus* and *Betula*) or are already so common in other areas that moderate planting efforts will have relatively small effects (*Quercus*). While *Gleditsia* was both a large producer of pollen and especially common as a street tree, its airborne pollen concentrations are generally low and it has been characterized as entomophilous (Schnabel and Hamrick 1995); moreover, it is rarely considered an important allergen. However, for *Platanus* × *acerifolia* historical street tree planting decisions have dramatically elevated pollen exposure, w ith large potential health consequences. Indeed, *Platanus* was predicted to be the second largest producer of allergenic pollen in NYC and was the fourth most collected pollen type. *Platanus* has also been planted widely in other urban areas around the world and is considered of substantial allergenic concern (Cariñanos et al. 2020; M. Nowak et al. 2012; Varela et al. 1994). Luckily, the relatively low numbers of *Platanus* × *acerifolia* in smaller size classes suggests that they are no longer being planted as frequently.

### Spatial variation and pollen dispersion

There is a growing body of literature demonstrating intra -municipal variation in airborne pollen concentrations (Katz et al. 2019, 2023; Katz and Batterman 2019; Weinberger et al. 2015, 2018; Zapata-Marin et al. 2021), due in part to spatial variation in plant abundance. For trees, there is evidence that trees have the strongest effects on airborne pollen concentrations at the neighborhood scale, i.e., within hundreds of meters to a few kilometers (Katz et al. 2019, 2023). This is relevant to the current study in part because the local nature of pollen dispersion cautions against using measurements from Lincoln Center as representative of the entire city. This could help to explain some of the mismatches between predicted pollen production and airborne pollen concentrations (even so, the top four genera for city-wide pollen production and airborne pollen measurements overlapped and both estimated pollen production and airborne pollen concentrations were highest for *Quercus*). Indeed the effects of local trees on airborne pollen concentration measurements were observed anecdotally; in the mid-2010s several large *Platanus* and *Morus* trees near Lincoln Center were removed during construction; subsequently, lower pollen concentrations were recorded for those genera (G. Robinson, *personal communication*). This leads to a critical point for tree planting decisions: even though certain taxa may not be especially important producers of pollen on an municipal scal e, they could still be of substantial local concern. Thus, a better understanding of what trees occur and how their pollen disperses is required for fully -informed tree planting decisions.

### Sex bias in tree planting

There has been considerable discussion of the disproportionate planting of male trees for diecious species in popular science (Ogren 2015). However, based on the data from this study, ‘botanical sexism’ is unlikely to play a major role in pollen exposure in NYC. Of the top pollen producing taxa for which we could estimate pollen production, the most i mportant diecious species belong to the *Morus* genus. However, very few individuals in this genus were street trees. Anecdotally, *Morus alba* trees are common as naturally-regenerating individuals, and are especially frequent along fence lines, vacant lots, and other less-managed areas, suggesting a minimal role for planting let alone botanical sexism in their establishment. In Baltimore, 0% of *Morus alba* trees were classified as planted, in Chicago, 25% of *Morus* trees were planted and in Hartford, CT 8% of *Morus rubra* trees were planted (D. J. Nowak 2012). The other species with varying sex ratios that we could estimate pollen production for (*Acer negundo, Acer rubrum*, and *Populus deltoides*) do not appear to be substantial producers of allergenic pollen across NYC; moreover, *Acer negundo* and *Populus deltoides* are early successional species and rarely planted intentionally. However, Cupressaceae does contain many diecious taxa, including several *Juniperus* species and was a moderate portion of airborne pollen (6% of airborne pollen), so we are unable to rule out its potential importance here. Furthermore, disproportionate planting of male trees of allergic concern could certainly be an issue in regions of the world where more of the species of allergenic concern are diecious (Cariñanos and Casares-Porcel 2011).

### Tree pollen allergenicity

Several sources describe the allergic potential of trees (Cariñanos et al. 2016; Ogren 2015; Pollen.com 2023), however the information in these metrics are sometimes contradictory between sources or based on expert opinion rather than observed epidemiological effects (Sousa-Silva et al. 2021). Thus, the connection between pollen exposure and health effects remains surprisingly poorly described. This may be due in part to the substantial measurement error in epidemiological analyses of allergenic pollen (Katz et al. 2019) that makes it challenging to effectively link pollen concentrations with health outcomes. This uncertainty, as well as the complicated nature of allergic sensitization, priming, and both intra - and inter-individual variability in dose -response relationships complicate efforts to connect pollen production to health outcomes. The shape of dose-response curves are directly relevant to tree management decisions such as tree planting. For example, if the effects of pollen on allergic responses are saturating, then attempts to reduce pollen may be wasted effort when there are already many nearby pollen sources. More importantly, allergenicity varies among taxa so the same concentrations of pollen could lead to substantially different health effects.

### Study limitations

The conclusions here are based upon data collected from 2009 – 2016; there were changes in tree composition both within that period and since then (Maxwell et al. 2021b). For example, *Fraxinus* trees have been decimated by the emerald ash borer (*Agrilus planipennis*); it is likely that *Fraxinus* pollen concentrations have dropped since the start of the airborne pollen collection period. However, large changes between the i-Tree survey in 2013 and the street tree survey in 2015 and 2016 are unlikely. A more important source of variation is expected to be the relatively low sample size of the i -Tree survey; for many taxa only a few individuals were recorded, which can have large effects on estimated pollen production. In the future, additional tree data from t he Urban Forest Inventory and Analysis will be available both in NYC and many other cities nationally (Edgar et al. 2021; US Forest Service 2016), which could help to address this concern.

Pollen production for several potentially important tree taxa remains unknown. The approach used here (i.e., applying equations for one or two species to all congenerics within *Quercus, Populus*, and *Ulmus*) is coarse and adds additional uncertainty to estimated total pollen production. These pollen production equations were also developed with street trees and for trees growing in generally open areas, so may not be as well suited for trees growing in natural areas, which generally experience higher competition. Thus, the results presented here likely overestimate total pollen production for species that are common in natural areas in NYC, including *Quercus*. This study also largely neglects the steps between pollen production and exposure; in addition to differential pollen production, different rates of entrainment and dispersion, such as the explosive release of pollen by *Morus* (Taylor et al. 2006), may also help to explain discrepancies in pollen production compared to airborne pollen measurements.

Another limitation of this study is that it focuses primarily on the potential allergenic consequences of planting street trees. This limitation is a practical consequence of the scant information available for trees growing on private land. The repeated decisions made by homeowners about what trees to plant – and what trees are available from nurseries – have strong influences on tree composition (Avolio et al. 2018); approximately 18% of trees in NYC are on properties of one or two family homes (Maxwell et al. 2021a). Tracking and understanding tree selection decisions by individuals is an area worthy of additional study and might help explain the abundance of certain taxa and perhaps offer a path towards solutions. For example, when ranking the importance of various ecosystem services and disservices provided by trees, “spreading pollen” was the 4^th^ lowest item out of 15 (Avolio et al. 2018), implying that greater visibility of this issue could help shift tree selection decisions for individuals or perhaps for the nurseries that supply trees to the public.

### Generalizability

Tree cover and composition varies substantially among and within cities and countries due to a range of historical and contemporary circumstances (Roman et al. 2018) as does their allergenic potential (D. J. Nowak and Ogren 2021). The results presented here are specific to NYC but may be somewhat similar to other cities in eastern North America (Sousa-Silva et al. 2021), as similar urban processes act to homogenize urban plant composition (Groffman et al. 2014). However, the potential effect of tree planting campaigns on pollen production will likely vary in part on the proportion of planted to unplanted trees, which varies dramatically within North America (D. J. Nowak 2012).

### Conclusion

There is substantial interest in tree planting campaigns (Eisenman et al. 2021b; Sousa-Silva et al. 2023) as cities turn towards green infrastructure to increase cooling and other ecosystem services provided by trees (Doroski et al. 2020; Rahman et al. 2020). However, to optimize the benefits of tree planting campaigns, ecosystem disservices such as allergenic pollen must also be taken into account (Eisenman et al. 2019). Urban foresters balance multiple suitability concerns including tree diversity and stock availability when selecting trees for planting; thus, considerations besides allergenicity often can and should determine tree selection decisions. However, the results presented here show how tree planting decisions have had long-term consequences for pollen production and exposure (i.e., for *Platanus* × *acerifolia*), providing strong incentive to include allergenic potential in upcoming tree selection decisions. Overall, the results from this study suggest that planting practices over the last several years are not substantially exacerbating citywide allergenic pollen exposure in NYC. Nonetheless, the increasing evidence of intra-municipal variation in airborne pollen concentrations suggests that planting allergenic pollen producing trees will have notable local effects, implying that the worst offending trees should not be planted in residential areas. From an allergy perspective, potential substitutes include the many taxa of trees that are insect-pollinated, have non-allergenic pollen, or are females of diecious species.

## Acknowledgements

I gratefully acknowledge Dr. Guy Robinson for providing airborne pollen concentration measurements and the New York City Department of Parks and Recreation for their role in collecting and providing tree-related data in NYC. I also thank EnergyBDDO for their interest in allergenic pollen production and for securing funding for this study.

## Supporting Information

**Table SI 1.** Extended version of Table 1

| genus | i-Tree<br>relative<br>number | i-Tree<br>relative<br>number | Street tree<br>relative<br>basal area | Street tree<br>relative<br>number | Pollen<br>mean<br>(grains/m<sup>3</sup>) | Pollen<br>max<br>(grains/m<sup>3</sup>) | Wind<br>pollinated | Of<br>allergenic<br>concern |
| --- | --- | --- | --- | --- | --- | --- | --- | --- |
| *Quercus* | 26.52 | 10.52 | 18.21 | 12.71 | 48.69088 | 2158.832 | + | + |
| *Acer* | 19.68 | 11.78 | 15.57 | 13.61 | 2.01005 | 119.3182 | + | + |
| *Platanus* | 8.29 | 2.38 | 34.47 | 13.34 | 24.20304 | 2727.623 | + | + |
| *Liquidambar* | 3.49 | 1.58 | 1.38 | 1.63 | 1.058717 | 87.20539 | + | ~ |
| *Tilia* | 3.22 | 0.96 | 5.05 | 7.86 | 0.106303 | 11.04798 | - | ~ |
| *Robinia* | 3.16 | 4.08 | 0.3 | 0.27 | 0.006208 | 4 | - | - |
| *Ulmus* | 2.15 | 1.19 | 1.95 | 2.29 | 2.328864 | 176.7677 | + | + |
| *Crataegus* | 2.11 | 1.25 | 0.05 | 0.51 | 0 | 0 | - | - |
| *Fagus* | 2.04 | 0.26 | 0.02 | 0.06 | 0.300397 | 27.00617 | + | ~ |
| *Fraxinus* | 1.78 | 0.57 | 3.12 | 3.06 | 4.009975 | 437.5 | + | + |
| *Picea* | 1.75 | 1.09 | 0.05 | 0.1 | NA | NA | + | - |
| *Betula* | 1.66 | 4.87 | 0.05 | 0.21 | 24.97922 | 2396.825 | + | + |
| *Pyrus* | 1.59 | 2.49 | 4.39 | 9.04 | NA | NA | - | - |
| *Tsuga* | 1.58 | 0.82 | 0 | 0.01 | 0.287978 | 33.14394 | + | - |
| *Gleditsia* | 1.52 | 0.92 | 6.11 | 9.85 | NA | NA | ~ | ~ |
| *Celtis* | 1.5 | 2.18 | 0.1 | 0.37 | 0.007357 | 2.209596 | + | + |
| *Unknown* | 1.42 | 3.4 | 0 | 0 | NA | NA |  |  |
| *Sophora* | 1.41 | 0.46 | 0 | 0 | NA | NA | - | - |
| *Thuja* | 1.1 | 6.32 | 0.01 | 0.05 | NA | NA | + | + |
| *Ailanthus* | 1.02 | 5.54 | 0.13 | 0.12 | NA | NA |  | ~ |
| *Prunus* | 0.93 | 3.35 | 1.37 | 6.39 | 0.1754 | 43.65079 | - | - |
| *Phellodendron* | 0.81 | 0.09 | 0.02 | 0.03 | NA | NA | + | ~ |
| *Maclura* | 0.79 | 0.1 | 0 | 0 | NA | NA | + | + |
| *Zelkova* | 0.71 | 0.72 | 1.91 | 4.49 | 0.424032 | 295.6349 | + |  |
| *Morus* | 0.69 | 0.82 | 0.29 | 0.18 | 45.94652 | 4342.593 | + | + |
| *Cedrus* | 0.68 | 2.54 | 0 | 0.02 | 0.019144 | 15 | + | - |
| *Malus* | 0.65 | 0.89 | 0.1 | 0.54 | NA | NA | - | - |
| *Sassafras* | 0.63 | 4.71 | 0.03 | 0.03 | NA | NA | - | - |
| *Ginkgo* | 0.57 | 0.53 | 1.61 | 3.22 | 1.503001 | 311.1111 | + | ~ |
| *Abies* | 0.53 | 0.17 | 0 | 0 | 0.022198 | 20 | + | - |
| *Cornus* | 0.5 | 1.78 | 0.09 | 0.49 | 0.006659 | 6 | - | - |
| *Taxus* | 0.49 | 1.78 | 0 | 0 | NA | NA | + | + |
| *Ficus* | 0.48 | 2.15 | 0 | 0 | NA | NA | - | - |
| *Amelanchier* | 0.46 | 0.47 | 0.03 | 0.31 | NA | NA | - | - |
| *Juniperus* | 0.46 | 1.2 | 0.04 | 0.17 | NA | NA | + | + |
| *Salix* | 0.45 | 0.46 | 0.07 | 0.04 | 0.963795 | 95.2381 |  |  |
| *Juglans* | 0.36 | 0.09 | 0.04 | 0.05 | 0.795508 | 150.2525 | + | + |
| *Morella* | 0.3 | 3.65 | 0 | 0 | NA | NA | - | - |
| *Pinus* | 0.26 | 0.43 | 0.08 | 0.11 | NA | NA | + | - |
| *Carya* | 0.24 | 2.84 | 0.02 | 0.02 | 1.105865 | 117.1086 | + | + |
| *Chamaecyparis* | 0.23 | 0.2 | 0.02 | 0.07 | NA | NA | + | + |
| *Ilex* | 0.2 | 0.67 | 0.01 | 0.02 | 0.015753 | 10 | - | - |
| *Rhododendron* | 0.18 | 0.64 | 0 | 0 | NA | NA | - | - |
| *Gymnocladus* | 0.17 | 0.09 | 0.09 | 0.52 | NA | NA | - | - |
| *Koelreuteria* | 0.17 | 0.29 | 0.09 | 0.57 | NA | NA | - | - |
| *Cephalotaxus* | 0.14 | 0.11 | 0 | 0 | NA | NA | + | ~ |
| *Hibiscus* | 0.14 | 0.83 | 0 | 0 | NA | NA | - | - |
| *Lagerstroemia* | 0.14 | 0.26 | 0.01 | 0.07 | NA | NA | - | - |
| *Magnolia* | 0.12 | 0.27 | 0.04 | 0.15 | NA | NA | - | - |
| *Liriodendron* | 0.09 | 1.32 | 0.27 | 0.16 | NA | NA | - | - |
| *Rhus* | 0.07 | 0.96 | 0 | 0 | 0.017938 | 8 | - | - |
| *Albizia* | 0.06 | 0.2 | 0.01 | 0.02 | NA | NA | - | - |
| *Nyssa* | 0.06 | 0.17 | 0.01 | 0.04 | 0.00424 | 2.209596 | - | - |
| *Aucuba* | 0.03 | 0.26 | 0 | 0 | NA | NA | - | - |
| *Diospyros* | 0.03 | 0.17 | 0 | 0 | NA | NA | - | - |
| *Photinia* | 0.03 | 0.09 | 0 | 0 | NA | NA | - | - |
| *Carpinus* | 0.02 | 0.11 | 0.1 | 0.62 | NA | NA | + | + |
| *Cercis* | 0.02 | 0.17 | 0.08 | 0.62 | NA | NA | - | - |
| *Ligustrum* | 0.02 | 0.4 | 0 | 0 | 0.002452 | 2.209596 | - | - |
| *Vitis* | 0.02 | 0.32 | 0 | 0 | NA | NA | ~ | - |
| *Aesculus* | 0.01 | 0.09 | 0.24 | 0.2 | NA | NA | - | - |
| *Buddleja* | 0.01 | 0.09 | 0 | 0 | NA | NA | - | - |
| *Castanea* | 0.01 | 0.09 | 0.01 | 0.03 | NA | NA | ~ | ~ |
| *Cryptomeria* | 0.01 | 0.09 | 0 | 0 | NA | NA | + | + |
| *Hydrangea* | 0.01 | 0.11 | 0 | 0 | NA | NA | - | - |
| *Lonicera* | 0.01 | 0.09 | 0 | 0 | NA | NA | - | - |
| *Syringa* | 0.01 | 0.19 | 0.08 | 0.77 | NA | NA | - | - |
| *Tabebuia* | 0.01 | 0.09 | 0 | 0 | NA | NA | - | - |
| *Viburnum* | 0.01 | 0.09 | 0 | 0 | NA | NA | - | - |
| *Lindera* | 0 | 0.09 | 0 | 0 | NA | NA | - | - |
| *Schizomeria* | 0 | 0.09 | 0 | 0 | NA | NA | - | - |
| *Styphnolobium* | 0 | 0 | 1.64 | 2.97 | NA | NA | - | - |
| *Catalpa* | 0 | 0 | 0.11 | 0.08 | NA | NA | - | - |
| *Populus* | 0 | 0 | 0.11 | 0.07 | 2.053207 | 156.7852 | + | + |
| *Metasequoia* | 0 | 0 | 0.11 | 0.46 | NA | NA | + | - |
| *Maackia* | 0 | 0 | 0.07 | 0.34 | NA | NA | - | - |
| *Ostrya* | 0 | 0 | 0.05 | 0.17 | NA | NA | + | + |
| *Paulownia* | 0 | 0 | 0.04 | 0.04 | NA | NA | - | - |
| *Taxodium* | 0 | 0 | 0.04 | 0.22 | NA | NA | + | + |
| *Cercidiphyllum* | 0 | 0 | 0.03 | 0.14 | NA | NA | - | - |
| *Eucommia* | 0 | 0 | 0.01 | 0.14 | NA | NA | + | ~ |
| *Cladrastis* | 0 | 0 | 0.01 | 0.07 | NA | NA | - | - |
| *Corylus* | 0 | 0 | 0.01 | 0.05 | 0.009559 | 4.419192 | + | + |
| *Chionanthus* | 0 | 0 | 0.01 | 0.04 | NA | NA | - | - |
| *Halesia* | 0 | 0 | 0.01 | 0.03 | NA | NA | - | - |
| *Styrax* | 0 | 0 | 0.01 | 0.06 | NA | NA | - | - |
| *Pseudotsuga* | 0 | 0 | 0 | 0.01 | 0 | 0 | + | - |
| *Larix* | 0 | 0 | 0 | 0.01 | 0.01421 | 9 | + | - |
| *Parrotia* | 0 | 0 | 0 | 0.04 | NA | NA | + | ~ |
| *Alnus* | 0 | 0 | 0 | 0.01 | 0.352962 | 31.74603 | + | + |
| *Cotinus* | 0 | 0 | 0 | 0.01 | NA | NA | - | - |

**Table SI 2:** Summary of 2015 street tree database by species. Only living trees were included. Trees with a genus listed but not a species were retained under just the genus listing.

| species | Total basal<br>area (m2) | Trees (n) | Basal area<br>(proportion) | Proportion<br>of all trees |
| --- | --- | --- | --- | --- |
| *Platanus x acerifolia* | 24109.7 | 87014 | 0.344694 | 0.133423 |
| *Quercus palustris* | 10422.5 | 53185 | 0.149009 | 0.081551 |
| *Gleditsia triacanthos* var. *inermis* | 4274.801 | 64263 | 0.061116 | 0.098537 |
| *Acer platanoides* | 4181.253 | 34189 | 0.059779 | 0.052424 |
| *Acer saccharinum* | 3376.41 | 12277 | 0.048272 | 0.018825 |
| *Pyrus calleryana* | 3072.143 | 58931 | 0.043922 | 0.090362 |
| *Tilia cordata* | 2165.223 | 29742 | 0.030956 | 0.045605 |
| *Fraxinus pennsylvanica* | 1770.462 | 16251 | 0.025312 | 0.024918 |
| *Acer rubrum* | 1400.903 | 17246 | 0.020029 | 0.026444 |
| *Zelkova serrata* | 1332.768 | 29258 | 0.019054 | 0.044863 |
| *Styphnolobium japonicum* | 1148.632 | 19338 | 0.016422 | 0.029652 |
| *Ginkgo biloba* | 1129.135 | 21024 | 0.016143 | 0.032237 |
| *Liquidambar styraciflua* | 965.2699 | 10657 | 0.0138 | 0.016341 |
| *Quercus rubra* | 964.9451 | 8400 | 0.013796 | 0.01288 |
| *Ulmus americana* | 936.3941 | 7975 | 0.013388 | 0.012228 |
| *Tilia americana* | 884.5466 | 13530 | 0.012646 | 0.020746 |
| *Prunus* | 723.8313 | 29279 | 0.010349 | 0.044895 |
| *Acer* | 644.2233 | 7080 | 0.00921 | 0.010856 |
| *Tilia tomentosa* | 481.166 | 7995 | 0.006879 | 0.012259 |
| *Quercus phellos* | 419.1621 | 3184 | 0.005993 | 0.004882 |
| *Acer platanoides* 'Crimson King' | 398.8446 | 5923 | 0.005702 | 0.009082 |
| *Acer pseudoplatanus* | 381.8622 | 2731 | 0.005459 | 0.004188 |
| *Fraxinus* | 295.4005 | 2609 | 0.004223 | 0.004001 |
| *Acer saccharum* | 255.8671 | 2844 | 0.003658 | 0.004361 |
| *Ulmus parvifolia* | 242.2366 | 5345 | 0.003463 | 0.008196 |
| *Quercus bicolor* | 219.9121 | 6598 | 0.003144 | 0.010117 |
| *Robinia pseudoacacia* | 208.1286 | 1784 | 0.002976 | 0.002735 |
| *Morus* | 202.9014 | 1157 | 0.002901 | 0.001774 |
| *Liriodendron tulipifera* | 189.3352 | 1076 | 0.002707 | 0.00165 |
| *Ulmus pumila* | 185.6489 | 1595 | 0.002654 | 0.002446 |
| *Aesculus hippocastanum* | 158.8608 | 1096 | 0.002271 | 0.001681 |
| *Quercus velutina* | 138.8934 | 1327 | 0.001986 | 0.002035 |
| *Quercus coccinea* | 137.3773 | 1465 | 0.001964 | 0.002246 |
| *Quercus shumardii* | 135.8896 | 1487 | 0.001943 | 0.00228 |
| *Fraxinus americana* | 116.6271 | 1121 | 0.001667 | 0.001719 |
| *Prunus cerasifera* | 114.9823 | 6879 | 0.001644 | 0.010548 |
| *Quercus alba* | 101.8875 | 1686 | 0.001457 | 0.002585 |
| *Acer campestre* | 91.05352 | 2550 | 0.001302 | 0.00391 |
| *Ailanthus altissima* | 88.79259 | 756 | 0.001269 | 0.001159 |
| *Acer ginnala* | 88.73685 | 2049 | 0.001269 | 0.003142 |
| *Catalpa* | 79.85678 | 551 | 0.001142 | 8.45E-04 |
| *Metasequoia glyptostroboides* | 77.92014 | 3020 | 0.001114 | 0.004631 |
| *Prunus virginiana* | 76.75927 | 4888 | 0.001097 | 0.007495 |
| *Quercus acutissima* | 74.87482 | 2244 | 0.00107 | 0.003441 |
| *Malus* | 73.1515 | 3527 | 0.001046 | 0.005408 |
| *Celtis occidentalis* | 73.04915 | 2382 | 0.001044 | 0.003652 |
| *Quercus robur* | 71.66634 | 1644 | 0.001025 | 0.002521 |
| *Koelreuteria paniculata* | 64.75331 | 3719 | 9.26E-04 | 0.005703 |
| *Gymnocladus dioicus* | 62.77614 | 3364 | 8.98E-04 | 0.005158 |
| *Populus deltoides* | 61.1643 | 276 | 8.74E-04 | 4.23E-04 |
| *Syringa reticulata* | 52.8563 | 4568 | 7.56E-04 | 0.007004 |
| *Cercis canadensis* | 52.75344 | 3801 | 7.54E-04 | 0.005828 |
| *Salix babylonica* | 50.50923 | 282 | 7.22E-04 | 4.32E-04 |
| *Maackia amurensis* | 48.28376 | 2197 | 6.90E-04 | 0.003369 |
| *Prunus serotina* | 42.8559 | 607 | 6.13E-04 | 9.31E-04 |
| *Acer palmatum* | 39.14223 | 1130 | 5.60E-04 | 0.001733 |
| *Ostrya virginiana* | 35.81569 | 1081 | 5.12E-04 | 0.001658 |
| *Crataegus* | 34.21601 | 2988 | 4.89E-04 | 0.004582 |
| *Cornus florida* | 32.64268 | 1552 | 4.67E-04 | 0.00238 |
| *Carpinus caroliniana* | 31.53856 | 1517 | 4.51E-04 | 0.002326 |
| *Juglans nigra* | 29.33539 | 295 | 4.19E-04 | 4.52E-04 |
| *Carpinus betulus* | 28.15324 | 2099 | 4.03E-04 | 0.003218 |
| *Juniperus virginiana* | 26.81198 | 1101 | 3.83E-04 | 0.001688 |
| *Paulownia tomentosa* | 26.35848 | 245 | 3.77E-04 | 3.76E-04 |
| *Quercus imbricaria* | 24.71319 | 1049 | 3.53E-04 | 0.001608 |
| *Taxodium distichum* | 23.64201 | 1261 | 3.38E-04 | 0.001934 |
| *Pinus strobus* | 22.91387 | 202 | 3.28E-04 | 3.10E-04 |
| *Picea abies* | 22.86928 | 355 | 3.27E-04 | 5.44E-04 |
| *Amelanchier* | 19.94963 | 2032 | 2.85E-04 | 0.003116 |
| *Cercidiphyllum japonicum* | 19.34512 | 911 | 2.77E-04 | 0.001397 |
| *Pinus* | 18.68083 | 289 | 2.67E-04 | 4.43E-04 |
| *Sassafras albidum* | 18.50145 | 171 | 2.65E-04 | 2.62E-04 |
| *Betula nigra* | 18.49436 | 565 | 2.64E-04 | 8.66E-04 |
| *Quercus macrocarpa* | 18.21871 | 515 | 2.60E-04 | 7.90E-04 |
| *Magnolia* | 17.24988 | 699 | 2.47E-04 | 0.001072 |
| *Carya glabra* | 16.25825 | 99 | 2.32E-04 | 1.52E-04 |
| *Cornus mas* | 16.25724 | 1066 | 2.32E-04 | 0.001635 |
| *Chamaecyparis thyoides* | 14.90585 | 355 | 2.13E-04 | 5.44E-04 |
| *Fagus grandifolia* | 13.08068 | 273 | 1.87E-04 | 4.19E-04 |
| *Phellodendron amurense* | 11.61123 | 183 | 1.66E-04 | 2.81E-04 |
| *Eucommia ulmoides* | 10.23704 | 915 | 1.46E-04 | 0.001403 |
| *Betula papyrifera* | 10.20613 | 535 | 1.46E-04 | 8.20E-04 |
| *Populus tremuloides* | 8.978878 | 83 | 1.28E-04 | 1.27E-04 |
| *Betula pendula* | 8.790382 | 300 | 1.26E-04 | 4.60E-04 |
| *Populus grandidentata* | 8.766567 | 94 | 1.25E-04 | 1.44E-04 |
| *Acer nigrum* | 8.396669 | 114 | 1.20E-04 | 1.75E-04 |
| *Thuja occidentalis* | 7.703492 | 328 | 1.10E-04 | 5.03E-04 |
| *Picea* | 7.64066 | 202 | 1.09E-04 | 3.10E-04 |
| *Pinus resinosa* | 7.347782 | 106 | 1.05E-04 | 1.63E-04 |
| *Cladrastis kentukea* | 7.185129 | 479 | 1.03E-04 | 7.34E-04 |
| *Corylus colurna* | 7.131924 | 317 | 1.02E-04 | 4.86E-04 |
| *Magnolia acuminata* | 6.971298 | 169 | 9.97E-05 | 2.59E-04 |
| *Carpinus japonica* | 6.849688 | 426 | 9.79E-05 | 6.53E-04 |
| *Aesculus x carnea* | 6.587719 | 116 | 9.42E-05 | 1.78E-04 |
| *Lagerstroemia* | 6.431146 | 442 | 9.19E-05 | 6.78E-04 |
| *Quercus falcata* | 6.42912 | 83 | 9.19E-05 | 1.27E-04 |
| *Cornus kousa* | 6.063783 | 302 | 8.67E-05 | 4.63E-04 |
| *Acer negundo* | 5.507416 | 64 | 7.87E-05 | 9.81E-05 |
| *Aesculus glabra* | 5.482081 | 75 | 7.84E-05 | 1.15E-04 |
| *Syringa pekinensis* | 5.475494 | 462 | 7.83E-05 | 7.08E-04 |
| *Castanea mollissima* | 5.131945 | 173 | 7.34E-05 | 2.65E-04 |
| *Picea pungens* | 5.042765 | 126 | 7.21E-05 | 1.93E-04 |
| *Albizia julibrissin* | 4.801571 | 163 | 6.86E-05 | 2.50E-04 |
| *Acer buergerianum* | 4.78941 | 110 | 6.85E-05 | 1.69E-04 |
| *Chionanthus retusus* | 4.62625 | 234 | 6.61E-05 | 3.59E-04 |
| *Cornus alternifolia* | 4.568485 | 280 | 6.53E-05 | 4.29E-04 |
| *Cercis reniformis* | 4.414446 | 219 | 6.31E-05 | 3.36E-04 |
| *Acer tataricum* | 4.392657 | 153 | 6.28E-05 | 2.35E-04 |
| *Fagus sylvatica* | 3.983237 | 125 | 5.69E-05 | 1.92E-04 |
| *Ilex* | 3.899123 | 138 | 5.57E-05 | 2.12E-04 |
| *Crataegus crusgalli* var. *inermis* | 3.886455 | 341 | 5.56E-05 | 5.23E-04 |
| *Nyssa sylvatica* | 3.855546 | 288 | 5.51E-05 | 4.42E-04 |
| *Halesia diptera* | 3.788661 | 221 | 5.42E-05 | 3.39E-04 |
| *Styrax japonicus* | 3.785114 | 392 | 5.41E-05 | 6.01E-04 |
| *Pinus rigida* | 3.428898 | 33 | 4.90E-05 | 5.06E-05 |
| *Acer truncatum* | 3.201385 | 59 | 4.58E-05 | 9.05E-05 |
| *Acer griseum* | 2.793992 | 220 | 3.99E-05 | 3.37E-04 |
| *Pseudotsuga menziesii* | 2.647553 | 85 | 3.79E-05 | 1.30E-04 |
| *Magnolia grandiflora* | 2.554825 | 132 | 3.65E-05 | 2.02E-04 |
| *Larix laricina* | 2.512768 | 46 | 3.59E-05 | 7.05E-05 |
| *Pinus nigra* | 2.511755 | 37 | 3.59E-05 | 5.67E-05 |
| *Cedrus atlantica* | 2.182394 | 87 | 3.12E-05 | 1.33E-04 |
| *Parrotia persica* | 2.068891 | 277 | 2.96E-05 | 4.25E-04 |
| *Pinus sylvestris* | 1.972617 | 25 | 2.82E-05 | 3.83E-05 |
| *Tsuga canadensis* | 1.89357 | 88 | 2.71E-05 | 1.35E-04 |
| *Alnus glutinosa* | 1.718756 | 47 | 2.46E-05 | 7.21E-05 |
| *Cotinus coggygria* | 1.585998 | 58 | 2.27E-05 | 8.89E-05 |
| *Maclura pomifera* | 1.138068 | 29 | 1.63E-05 | 4.45E-05 |
| *Cedrus deodara* | 1.128947 | 72 | 1.61E-05 | 1.10E-04 |
| *Taxodium ascendens* | 1.032166 | 181 | 1.48E-05 | 2.78E-04 |
| *Chamaecyparis pisifera* | 1.011897 | 108 | 1.45E-05 | 1.66E-04 |
| *Pinus virginiana* | 0.113503 | 10 | 1.62E-06 | 1.53E-05 |

**Table SI 3:** Summary of 2013 i -Tree data by species.

| Genus | Species | Common Name | Trees | Trees percent | Basal Area |
| --- | --- | --- | --- | --- | --- |
| *Acer* | *platanoides* | Norway maple | 425,000 | 6.1 | 451,580 |
| *Quercus* | *palustris* | Pin oak | 123,000 | 1.8 | 352,640 |
| *Platanus* | *hybrida* | London planetree | 166,000 | 2.4 | 307,870 |
| *Quercus* | *alba* | White oak | 302,000 | 4.3 | 178,960 |
| *Quercus* | *velutina* | Black oak | 61,000 | 0.9 | 134,720 |
| *Acer* | *saccharinum* | Silver maple | 64,000 | 0.9 | 130,000 |
| *Liquidambar* | *styraciflua* | Sweetgum | 110,000 | 1.6 | 129,610 |
| *Acer* | *rubrum* | Red maple | 230,000 | 3.3 | 119,280 |
| *Robinia* | *pseudoacacia* | Black locust | 285,000 | 4.1 | 117,550 |
| *Quercus* | *rubra* | Northern red oak | 98,000 | 1.4 | 112,590 |
| *Quercus* | *species* | Oak spp | 36,000 | 0.5 | 97,820 |
| *Quercus* | *bicolor* | Swamp white oak | 89,000 | 1.3 | 86,810 |
| *Crataegus* | *phaenopyrum* | Washington hawthorn | 34,000 | 0.5 | 73,990 |
| *Fagus* | *sylvatica* | Copper beech | 6,000 | 0.1 | 64,660 |
|  | *'purpurea'* |  |  |  |  |
| *Picea* | *abies* | Norway spruce | 51,000 | 0.7 | 61,040 |
| *Tsuga* | *canadensis* | Eastern hemlock | 51,000 | 0.7 | 58,220 |
| *Gleditsia* | *triacanthos* | Honeylocust | 64,000 | 0.9 | 56,600 |
| *Tilia* | *cordata* | Littleleaf linden | 27,000 | 0.4 | 54,900 |
| *Pyrus* | *calleryana* | Callery pear | 160,000 | 2.3 | 54,820 |
| *Betula* | *lenta* | Black birch | 274,000 | 3.9 | 53,700 |
| *Unknown* | *species* | Hardwood | 237,000 | 3.4 | 52,720 |
| *Sophora* | *japonica* | Japanese pagoda tree | 32,000 | 0.5 | 52,350 |
| *Tilia* | *tomentosa* | Silver linden | 14,000 | 0.2 | 49,760 |
| *Ulmus* | *americana* | American elm | 21,000 | 0.3 | 47,200 |
| *Thuja* | *occidentalis* | Northern white-cedar | 395,000 | 5.7 | 38,810 |
| *Ailanthus* | *altissima* | Tree-of-heaven | 387,000 | 5.5 | 37,960 |
| *Fraxinus* | *americana* | White ash | 6,000 | 0.1 | 36,370 |
| *Ulmus* | *parvifolia* | Chinese elm | 48,000 | 0.7 | 31,710 |
| *Celtis* | *occidentalis* | Northern hackberry | 121,000 | 1.7 | 31,110 |
| *Phellodendron* | *amurense* | Amur corktree | 6,000 | 0.1 | 30,060 |
| *Maclura* | *pomifera* | Osage orange | 7,000 | 0.1 | 29,410 |
| *Zelkova* | *serrata* | Japanese zelkova | 50,000 | 0.7 | 26,300 |
| *Fraxinus* | *pennsylvanica* | Green ash | 26,000 | 0.4 | 26,240 |
| *Morus* | *species* | Mulberry spp | 57,000 | 0.8 | 25,580 |
| *Celtis* | *species* | Hackberry spp | 31,000 | 0.5 | 24,700 |
| *Cedrus* | *species* | Cedar spp | 165,000 | 2.4 | 24,300 |
| *Sassafras* | *albidum* | Sassafras | 329,000 | 4.7 | 23,560 |
| *Acer* | *palmatum* | Japanese maple | 38,000 | 0.5 | 22,060 |
| *Ginkgo* | *biloba* | Ginkgo | 37,000 | 0.5 | 21,130 |
| *Quercus* | *phellos* | Willow oak | 12,000 | 0.2 | 19,370 |
| *Abies* | *grandis* | Grand fir | 6,000 | 0.1 | 19,240 |
| *Amelanchier* | *species* | Serviceberry spp | 33,000 | 0.5 | 17,040 |
| *Taxus* | *species* | Yew spp | 110,000 | 1.6 | 15,760 |
| *Juglans* | *species* | Walnut spp | 6,000 | 0.1 | 13,360 |
| *Malus* | *spectabilis* | Asiatic apple | 7,000 | 0.1 | 13,070 |
| *Tilia* | *americana* | American basswood | 20,000 | 0.3 | 11,760 |
| *Morella* | *pensylvanica* | Bayberry | 255,000 | 3.6 | 11,200 |
| *Ficus* | *species* | Fig spp | 76,000 | 1.1 | 11,170 |
| *Fagus* | *species* | Beech spp | 12,000 | 0.2 | 11,090 |
| *Juniperus* | *virginiana* | Eastern red cedar | 44,000 | 0.6 | 10,840 |
| *Cornus* | *florida* | Flowering dogwood | 73,000 | 1.0 | 9,480 |
| *Salix* | *fragilis* | Crack willow | 8,000 | 0.1 | 9,470 |
| *Cornus* | *species* | Dogwood spp | 45,000 | 0.7 | 8,850 |
| *Pinus* | *species* | Pine spp | 18,000 | 0.3 | 8,640 |
| *Prunus* | *serotina* | Black cherry | 25,000 | 0.4 | 8,490 |
| *Chamaecyparis* | *thyoides* | Atlantic white cedar | 14,000 | 0.2 | 8,440 |
| *Prunus* | *species* | Plum spp | 86,000 | 1.2 | 8,090 |
| *Malus* | *species* | Apple spp | 43,000 | 0.6 | 7,570 |
| *Ilex* | *aquifolium* | English holly | 35,000 | 0.5 | 6,860 |
| *Salix* | *species* | Willow spp | 18,000 | 0.3 | 6,680 |
| *Ficus* | *carica* | Common fig | 74,000 | 1.1 | 6,540 |
| *Koelreuteria* | *paniculata* | Goldenrain tree | 20,000 | 0.3 | 6,390 |
| *Gymnocladus* | *dioicus* | Kentucky coffeetree | 6,000 | 0.1 | 6,280 |
| *Rhododendron* | *azalea* | Azalea | 31,000 | 0.4 | 5,850 |
| *Prunus* | *pissardii* | Purpleleaf plum | 29,000 | 0.4 | 5,450 |
| *Hibiscus* | *syriacus* | Rose-of-sharon | 58,000 | 0.8 | 5,210 |
| *Cephalotaxus* | *fortunei* | Chinese plum yew | 8,000 | 0.1 | 5,090 |
| *Lagerstroemia* | *indica* | Common crapemyrtle | 18,000 | 0.3 | 5,060 |
| *Juniperus* | *virginiana silicicol* | Southern redcedar | 12,000 | 0.2 | 4,650 |
| *Magnolia* | *species* | Magnolia spp | 19,000 | 0.3 | 4,470 |
| *Pyrus* | *species* | Pear spp | 14,000 | 0.2 | 4,340 |
| *Crataegus* | *species* | Hawthorn spp | 46,000 | 0.7 | 4,080 |
| *Prunus* | *serrulata* | Kwanzan cherry | 30,000 | 0.4 | 4,080 |
| *Acer* | *pseudoplatanus* | Sycamore maple | 6,000 | 0.1 | 4,040 |
| *Carya* | *glabra* | Pignut hickory | 159,000 | 2.3 | 3,750 |
| *Picea* | *pungens* | Blue spruce | 19,000 | 0.3 | 3,710 |
| *Carya* | *alba* | Mockernut hickory | 18,000 | 0.3 | 3,690 |
| *Malus* | *tschonoskii* | Crabapple | 12,000 | 0.2 | 3,640 |
| *Liriodendron* | *tulipifera* | Tulip tree | 92,000 | 1.3 | 3,520 |
| *Fraxinus* | *nigra* | Black ash | 8,000 | 0.1 | 3,410 |
| *Tilia* | *x vulgaris* | Common linden | 6,000 | 0.1 | 3,340 |
| *Prunus* | *persica* | Peach | 32,000 | 0.5 | 3,260 |
| *Betula* | *occidentalis* | Water birch | 14,000 | 0.2 | 2,740 |
| *Taxus* | *canadensis* | Canada yew | 14,000 | 0.2 | 2,600 |
| *Nyssa* | *sylvatica* | Black tupelo | 12,000 | 0.2 | 2,370 |
| *Albizia* | *species* | Albizia spp | 14,000 | 0.2 | 2,360 |
| *Rhus* | *species* | Sumac spp | 52,000 | 0.7 | 2,360 |
| *Thuja* | *species* | Red cedar spp | 46,000 | 0.7 | 2,060 |
| *Quercus* | *prinus* | Chestnut oak | 6,000 | 0.1 | 2,050 |
| *Acer* | *buergerianum* | Trident maple | 19,000 | 0.3 | 1,740 |
| *Juniperus* | *species* | Juniper spp | 28,000 | 0.4 | 1,720 |
| *Betula* | *populifolia* | Gray birch | 12,000 | 0.2 | 1,670 |
| *Betula* | *papyrifera* | Paper birch | 12,000 | 0.2 | 1,600 |
| *Prunus* | *x orthosepala* | Prunus | 6,000 | 0.1 | 1,570 |
| *Betula* | *nigra* | River birch | 22,000 | 0.3 | 1,360 |
| *Prunus* | *cerasus* | Sour cherry | 6,000 | 0.1 | 1,200 |
| *Prunus* | *subhirtella* | Higan cherry | 6,000 | 0.1 | 1,200 |
| *Photinia* | *species* | Chokeberry spp | 6,000 | 0.1 | 1,150 |
| *Aucuba* | *japonica* | Japanese aucuba | 18,000 | 0.3 | 1,140 |
| *Prunus* | *mume* | Japanese apricot | 8,000 | 0.1 | 1,050 |
| *Carya* | *x ludoviciana* | Hickory | 15,000 | 0.2 | 1,010 |
| *Cedrus* | *atlantica glauca* | Blue atlas cedar | 12,000 | 0.2 | 960 |
| *Ulmus* | *species* | Elm spp | 14,000 | 0.2 | 960 |
| *Diospyros* | *virginiana* | Common persimmon | 12,000 | 0.2 | 930 |
| *Pinus* | *strobus* | Eastern white pine | 12,000 | 0.2 | 930 |
| *Rhododendron* | *arborescens* | Smooth azalea | 14,000 | 0.2 | 900 |
| *Acer* | *negundo* | Boxelder | 6,000 | 0.1 | 840 |
| *Ligustrum* | *species* | Privet spp | 28,000 | 0.4 | 820 |
| *Acer* | *campestre* | Hedge maple | 6,000 | 0.1 | 800 |
| *Acer* | *species* | Maple spp | 22,000 | 0.3 | 690 |
| *Vitis* | *species* | Grape spp | 22,000 | 0.3 | 680 |
| *Carpinus* | *caroliniana* | American hornbeam | 8,000 | 0.1 | 670 |
| *Cercis* | *reniformis* | Southwestern redbud | 12,000 | 0.2 | 600 |
| *Betula* | *species* | Birch spp | 6,000 | 0.1 | 530 |
| *Lonicera* | *japonica* | Hall's honeysuckle | 6,000 | 0.1 | 530 |
| *Tabebuia* | *heterophylla* | White cedar | 6,000 | 0.1 | 530 |
| *Buddleja* | *species* | Butterflybush spp | 6,000 | 0.1 | 510 |
| *Salix* | *x sepulcralis* | Weeping willow | 6,000 | 0.1 | 510 |
|  | *simonk* |  |  |  |  |
| *Tsuga* | *species* | Hemlock spp | 6,000 | 0.1 | 510 |
| *Ilex* | *species* | Holly spp | 12,000 | 0.2 | 430 |
| *Hydrangea* | *species* | Hydrangea spp | 8,000 | 0.1 | 380 |
| *Quercus* | *fastigiata* | Fastigate pin oak | 7,000 | 0.1 | 360 |
| *Syringa* | *reticulata* | Japanese tree lilac | 7,000 | 0.1 | 340 |
| *Abies* | *fraseri* | Fraser fir | 6,000 | 0.1 | 300 |
| *Aesculus* | *hippocastanum* | Horse chestnut | 6,000 | 0.1 | 300 |
| *Castanea* | *dentata* | American chestnut | 6,000 | 0.1 | 300 |
| *Cryptomeria* | *species* | Japanese cedar spp | 6,000 | 0.1 | 300 |
| *Picea* | *species* | Spruce spp | 6,000 | 0.1 | 300 |
| *Carya* | *species* | Hickory spp | 6,000 | 0.1 | 290 |
| *Cornus* | *mas* | Cornelian cherry | 6,000 | 0.1 | 290 |
| *Prunus* | *yedoensis* | Yoshino flowering | 6,000 | 0.1 | 290 |
|  |  | cherry |  |  |  |
| *Viburnum* | *species* | Viburnum spp | 6,000 | 0.1 | 290 |
| *Rhus* | *hirta* | Staghorn sumac | 15,000 | 0.2 | 200 |
| *Crataegus* | *mollis* | Downy hawthorn | 7,000 | 0.1 | 150 |
| *Acer* | *saccharum* | Sugar maple | 6,000 | 0.1 | 130 |
| *Lindera* | *benzoin* | Spicebush | 6,000 | 0.1 | 130 |
| *Schizomeria* | *ovata* | Cherry birch | 6,000 | 0.1 | 130 |
| *Syringa* | *species* | Lilac spp | 6,000 | 0.1 | 30 |

